# Multidimensional Analysis of the Adult Human Heart in Health and Disease using Hierarchical Phase-Contrast Tomography (HiP-CT)

**DOI:** 10.1101/2023.10.09.561474

**Authors:** J. Brunet, A. C. Cook, C. L. Walsh, J. Cranley, P. Tafforeau, K. Engel, C. Berruyer, E. Burke O’Leary, A. Bellier, R. Torii, C. Werlein, D. D. Jonigk, M. Ackermann, K. Dollman, P. D. Lee

## Abstract

Cardiovascular diseases (CVDs) are a leading cause of death worldwide. Current clinical imaging modalities provide resolution adequate for diagnosis but are unable to provide detail of structural changes in the heart, across length-scales, necessary for understanding underlying pathophysiology of disease. Hierarchical Phase-Contrast Tomography (HiP-CT), using new (4^th^) generation synchrotron sources, potentially overcomes this limitation, allowing micron resolution imaging of intact adult organs with unprecedented detail. In this proof of principle study (n=2), we show the utility of HiP-CT to image whole adult human hearts ex-vivo: one ‘control’ without known cardiac disease and one with multiple known cardiopulmonary pathologies. The resulting multiscale imaging was able to demonstrate exemplars of anatomy in each cardiac segment along with novel findings in the cardiac conduction system, from gross (20 um/voxel) to cellular scale (2.2 um/voxel), non-destructively, thereby bridging the gap between macroscopic and microscopic investigations. We propose that the technique represents a significant step in virtual autopsy methods for studying structural heart disease, facilitating research into abnormalities across scales and age-groups. It opens up possibilities for understanding and treating disease; and provides a cardiac ‘blueprint’ with potential for in-silico simulation, device design, virtual surgical training, and bioengineered heart in the future.

## Introduction

Cardiovascular diseases (CVDs) are the leading cause of mortality worldwide, with estimated annual costs of €210, and $320 billion in Europe(1) the United States(2) respectively. However, the development of new treatments for CVD is often impaired by lack of anatomic imaging tools which can bridge between macro and micro scales. The heart is a complex multiscale muscular pump with interconnected components at both micro- and macro scales which need to work together as one functionally unit and which change during evolution of CVDs. Despite decades of anatomic description, important research questions remain including identifying the anatomic substrates for cardiac arrhythmia, mitral annular disjunction(3,4), atrial and ventricular mural architecture(5), as well as the changes that occur throughout life in the increasing number of patients living with congenital heart disease.

Imaging of cardiac anatomy has classically been conducted via dissection, supplemented by histologic sampling, and/or combined with light or electron microscopy(6),(7). While these provide information down to sub-cellular resolution, physical sectioning is destructive, labour intensive, and causes morphological alteration(8). Optical microscopy of tissue blocks is often limited by light penetration(9),(10). Optical clearing with light-sheet fluorescent microscopy (LSFM) is a promising approach, which has been applied to intact whole human hearts(11,12), but clearing is slow, expensive, and the majority of light-sheet microscopes cannot image organs of this size(11).

Clinically, X-ray computed tomography (CT), Magnetic Resonance Imaging (MRI), and Ultrasound (US), are used widely and allow for rapid investigation of patients at the macro-scale with identification of indicators of CVDs(13). However, the limited resolution of these modalities hinders correlation with micro-anatomic changes in disease. Even high-spatial resolution MRI, ex-vivo, can provide information only to 50 µm/voxel in rat heart(14), while ultrasound, achieves resolutions of around 100 µm(15,16)

Laboratory micro-CT (µCT) is able to achieve whole mouse or human fetal heart imaging near to cellular resolution, but with application of contrast agents(17,18). Nano-tomography can achieve sub-µm resolution(19), but the trade-off between field of view and resolution restricts sample size significantly(5). These limitations can potentially be addressed by synchrotron X-ray phase-contrast imaging (XPCI, also called tomography, XPCT, phase-contrast imaging, PCI, synchrotron computed tomography, sCT, and other acronyms). The brightness of synchrotron sources eliminates the need for contrast agents and provides sufficient resolution to resolve cardiac microstructure(5). Whole human fetal and small animal hearts have been imaged with XPCI down to 5.2 µm(20). Moreover, hierarchical techniques such as propagation-based XPCI(21) have been used to image regions of interest in whole rat hearts down to 0.65 µm^3^/voxel. Thus, XPCI enables multiscale structural analysis of 3D cardiac anatomy including cardiomyocyte architecture and remodeling. Nonetheless, imaging whole adult human hearts at high-resolution, while overcoming limitations in field of view and resolution, has been challenging.

Recently, we utilised the new 4^th^ generation synchrotron source at the European Synchrotron Radiation Facility (ESRF), the Extremely Brilliant Source (EBS), to develop an XPCI technique: Hierarchical Phase-Contrast Tomography (HiP-CT), capable of imaging intact adult human organs with cellular resolution(22).

In this proof of principle study, we demonstrate the remarkable capability of HiP-CT to explore, non-destructively, the micro-structural architecture of normal and structurally abnormal adult human hearts, ex-vivo, with unprecedented detail by focusing on exemplars of anatomy. We obtained whole heart image volumes with isotropic voxels of 20 µm, and subsequent regions of interest down to 6.5-2.2 µm/voxel without the need for sectioning. For the first time, this multi-scale technique bridges the gap between gross anatomy and histologic observations, enabling the identification and understanding of abnormalities from macro-, to micro-level without the use of contrast agents such as Iodine or resorting to organ-damaging procedures. We believe this represents a step-change in CT-based virtual autopsy techniques for studying the heart, with exciting new possibilities for understanding normal and diseased anatomy.

## Materials and Methods

### Sample preparation

Two hearts were obtained from bodies donated to the Laboratoire d’Anatomie des Alpes Françaises (LADAF) following the current French legislation for body donation; one control and one heart with multiple pathologies (Table 1). Written informed consent was obtained antemortem from all the donors. All dissections respected the memory of the deceased. Transport and imaging protocols were approved by the Health Research Authority and Integrated Research Application System (HRA and IRAS) (200429) and the French Health Ministry.

**Table. 1:**
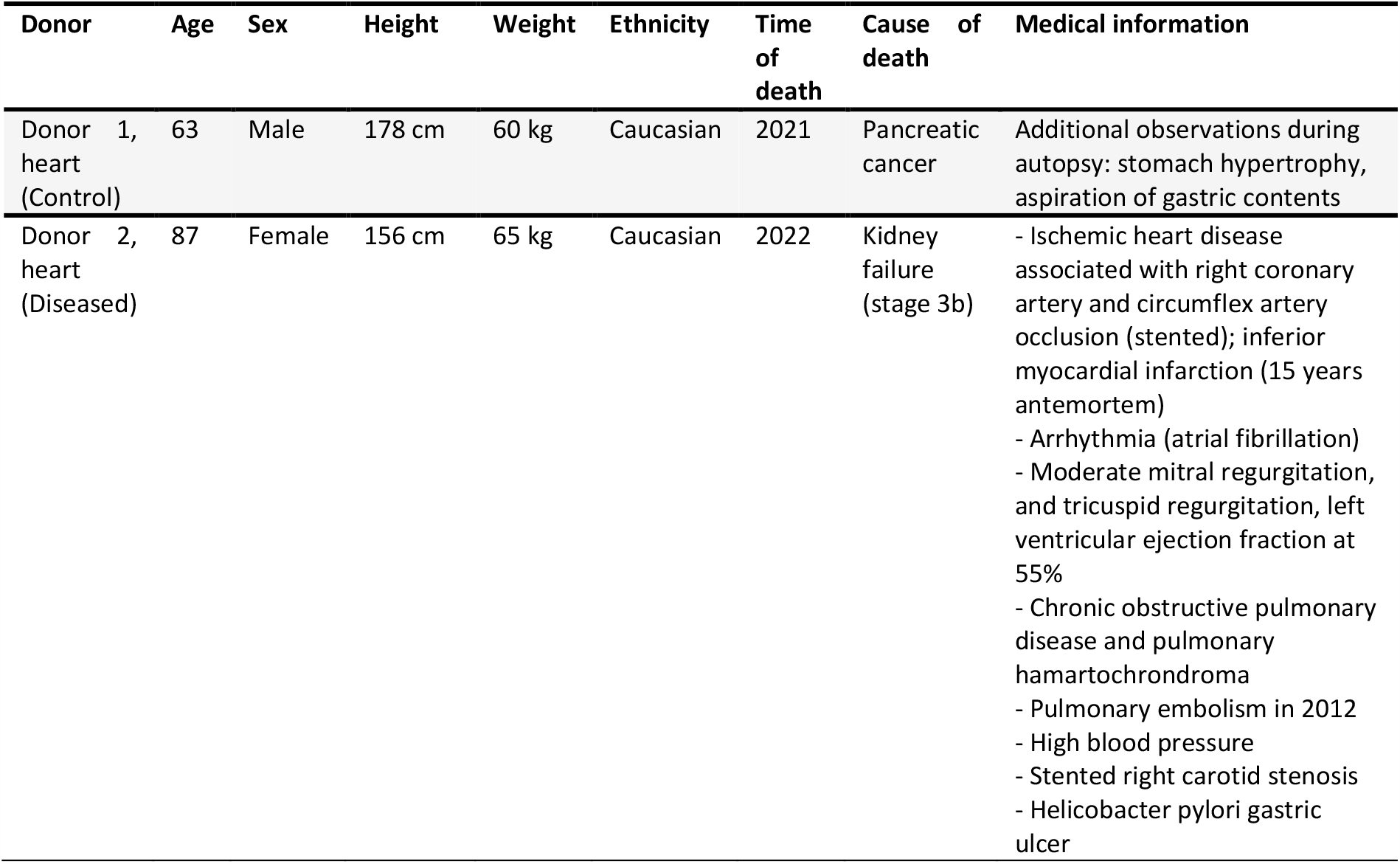
Medical information of the organ donors.

| Donor | Age | Sex | Height | Weight | Ethnicity | Time of death | Cause of death | Medical information |
| --- | --- | --- | --- | --- | --- | --- | --- | --- |
| Donor 1, heart (Control) | 63 | Male | 178 cm | 60 kg | Caucasian | 2021 | Pancreatic cancer | Additional observations during autopsy: stomach hypertrophy, aspiration of gastric contents |
| Donor 2, heart (Diseased) | 87 | Female | 156 cm | 65 kg | Caucasian | 2022 | Kidney failure (stage 3b) | - Ischemic heart disease associated with right coronary artery and circumflex artery occlusion (stented); inferior myocardial infarction (15 years antemortem)<br>- Arrhythmia (atrial fibrillation)<br>- Moderate mitral regurgitation, and tricuspid regurgitation, left ventricular ejection fraction at 55%<br>- Chronic obstructive pulmonary disease and pulmonary hamartochondroma<br>- Pulmonary embolism in 2012<br>- High blood pressure<br>- Stented right carotid stenosis<br>- Helicobacter pylori gastric ulcer |

Bodies were embalmed by injecting formalin through the right carotid artery after death, before storage at 3.6 °C. All dissections of the hearts were performed (abiding by COVID restrictions) between 16 April 2021 and 18 November 2021 at LADAF. During evisceration, the pericardium and lungs were removed. The hearts were prepared following the protocol of Brunet et al. (23). Briefly, hearts were post-fixed in 4% neutral buffered formalin for 4 days at room temperature, they were then partially dehydrated to 70% ethanol using multiple baths of increasing ethanol concentration (4 days per bath). A degassing cycle was performed at each step with a diaphragm vacuum pump (Vacuubrand, MV2) to avoid bubble formation during imaging. The hearts were stabilised in agar-ethanol gel with the cardiac apex facing downwards. Multiple steps of degassing were performed during the stabilisation process to remove any trapped air-bubbles. Subsequently, the container was sealed and stored until scanning.

### Propagation-based Synchrotron X-ray Tomography

Hearts were imaged on beamline BM18 at ESRF using HiP-CT as previously described by Walsh et al.(22). Scanning was performed using a quasi-parallel polychromatic beam tuned using a combination of molybdenum attenuators, with an average energy ranging between 94 and 126 keV. A schematic of the experimental setup is shown in Supplementary Fig. 1. Full-field tomography of the complete heart was performed with an isotropic voxel size of 20 µm, followed by local tomographic zooms in regions of interest (at 6.5um and 2.2 µm/voxel). Chosen regions of interest included: the atrial walls and left atrial appendage; the atrioventricular and arterial valves, the ventricular myocardium; and the coronary vasculature. Further zooms focussing on aspects of the cardiac conduction system, the sino-atrial node (SAN) and atrioventricular node (AVN), were selected by identifying and following the course of the respective nodal arteries on the low-resolution scans. Additional information on the method can be found in Supplementary materials. Comprehensive details regarding the scan parameters for the three distinct voxel sizes, including the scan time, number of projections, and X-ray beam energy, are provided in Supplementary Table 1.

### Data Processing and Analysis

Scans were reconstructed using a filtered back-projection algorithm, coupled with single-distance phase retrieval(24) and a 2D unsharp mask using PyHST2(25). Following vertical concatenation of scans, ring artefact correction was performed on the reconstructed slices using an updated Lyckegaard et al.(26) algorithm. Further details on the reconstruction pipeline can be found at Xian et al.(27). Data ranged from 150-915 GB per volume and required ultra-high-performance computing to process and visualise as described by Brunet et al.(23). Data was made publicly accessible through the web interface Neuroglancer, as detailed in the supplementary methods, and illustrated in Supplementary Fig. 2.

Volumes were manually registered for visualisation and semi-automatically segmented using a bounded region growing algorithm in VGSTUDIO MAX 3.5.1. For demonstration and comparison of normal and abnormal anatomy, datasets were resliced in the three orthogonal planes of the heart, namely 4-chamber, long axis, and short axis planes, as used in clinical imaging. 3D anatomic renderings were subsequently created using Cinematic rendering software (Siemens Healthineers)(28).

### Analysis

The thicknesses of the atrial wall and its constituent sub-layers (endocardium, myocardium, and epicardium) were manually measured at multiple evenly distributed locations (N=15). Differences in thickness in both control and diseased hearts, and in right vs left atrium were compared by Student’s t-test. Normality of data was assessed with a Shapiro-Wilk test. Assessment of the orientation of aggregates of myocytes in a mid-ventricular slice of the control heart, and of 3D mapping of the extent of mitral annular disjunction were performed as described in Supplemental Methods.

## Results

### Macroscopic normal and abnormal anatomy

HiP-CT successfully captured gross anatomy of the control and pathological intact adult human hearts at high resolution (approximately 20 × 20 × 20 µm^3^ voxel size) and was easily able to demonstrate the typical morphological features of the three major cardiac segments: the atria, ventricles, and arterial trunks.

As shown in Figs. 1-5 the phase-contrast provided by HiP-CT was able to depict subtle differences in soft tissue density in all tissue types of the heart including adipose, muscle, valvar and fibrous tissues. Consequently, in the diseased heart, we were able to make the post-mortem diagnosis of lipomatous hypertrophy of the interatrial ‘septum’ with the typical radiologic ‘hourglass’ configuration (Fig 2). Specifically, we were able to determine that excessive adipose tissue, continuous with epicardial fat, was sandwiched within the interatrial fold (Waterston’s or Sondergard’s groove) in between two thin layers of atrial myocardium, rather than being fatty infiltration of a ‘secondary septal’ structure (Fig 2b). We were also able to detect the presence of a heterogeneously composed thrombus located within the inter-atrial septum, trapped within, and protruding through a narrow patent foramen ovale (Fig. 2b,c). Again, this was an additional virtual autopsy diagnosis.

**Fig. 1:**
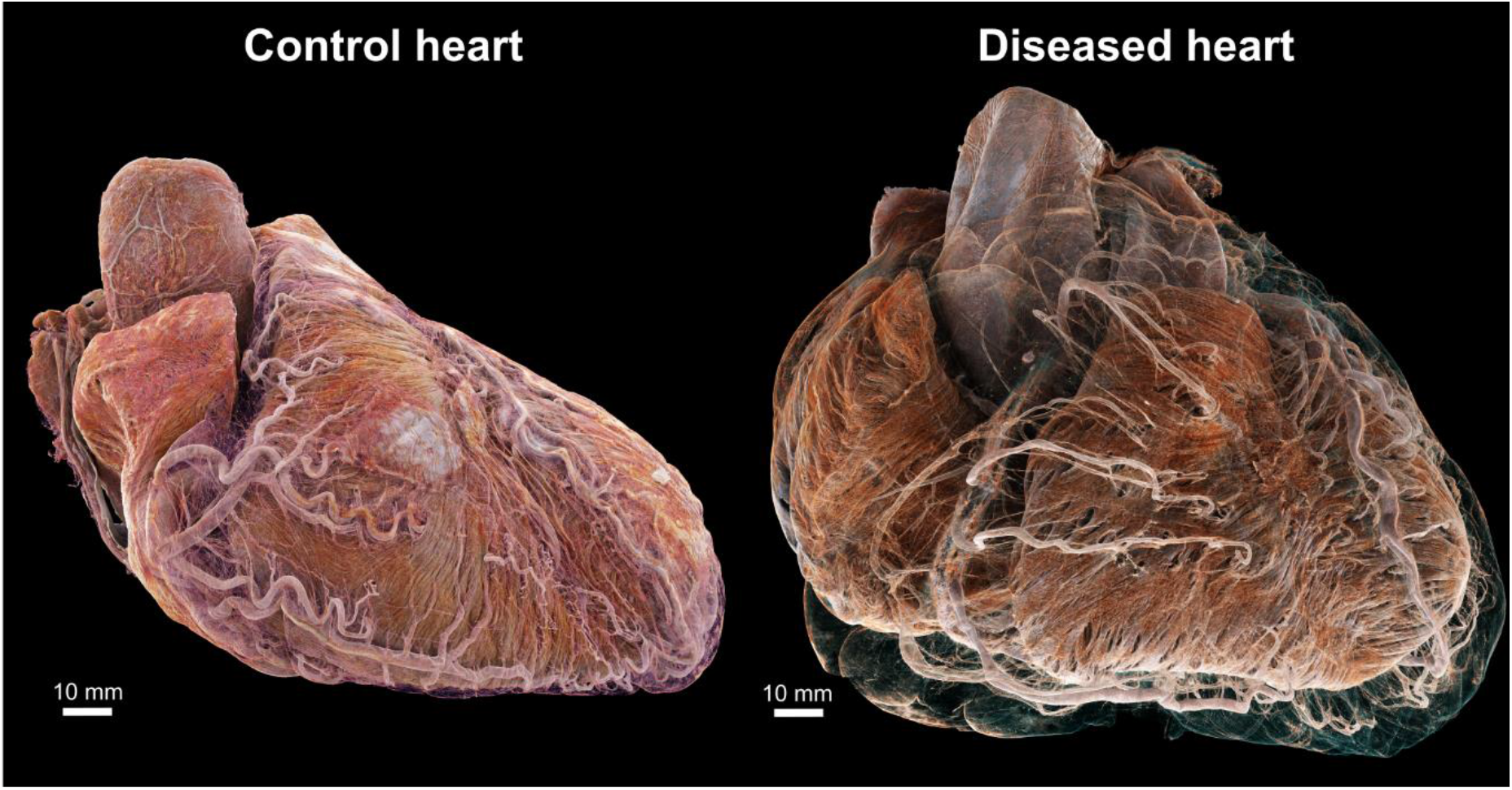
3D cinematic renderings of the normal (left) and abnormal (right) hearts in attitudinal position, i.e., as they would sit in the chest. The epicardial fat has been removed digitally to show the different course of the coronary vasculature between the two hearts. In the control, the coronary vasculature sits relatively close to the epicardial surface while in the disease heart, it is distant from the epicardium due to a thick layer of epicardial fat, while still being ‘anchored’ to the myocardium by smaller penetrating arteries. This gives rise to an unusual spiral configuration of the coronary arteries (see also Fig. 6g)

**Figure 2:**
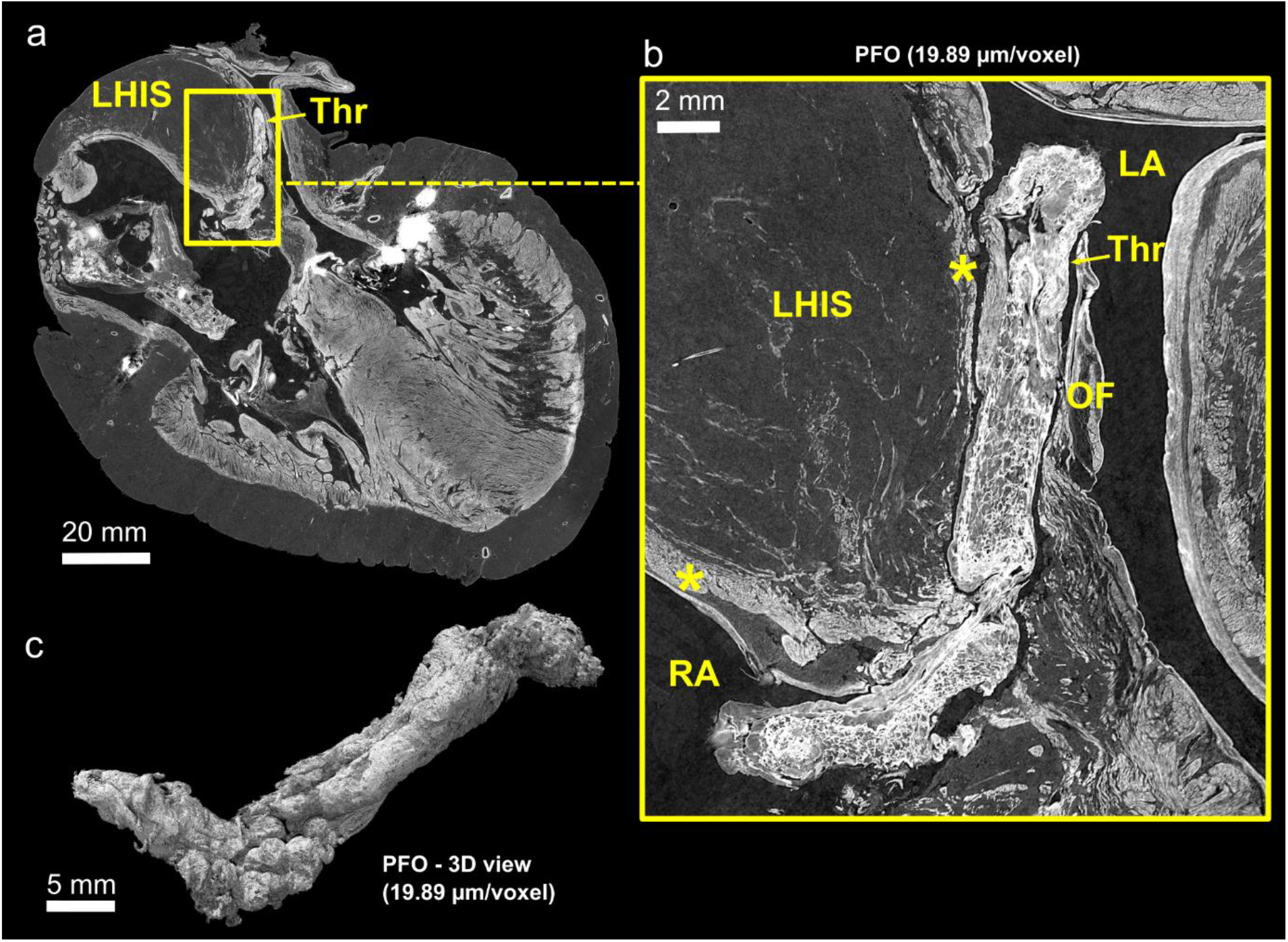
**(a)**, Cross section of thrombus protruding through a patent foramen ovale in the diseased heart at 20 µm/voxel. The characteristic ‘hourglass-shaped’ due to the lipomatous hypertrophy of the atrial septum (LHAS) can be observed. **(b)**, Digital zoom centred on the thrombus better showing its heterogeneous structure, and showing extra-cardiac fat distending the inter-atrial groove between two thin layers of atrial myocardium (yellow stars). **(c)**, 3D view of the thrombus after segmentation. The cylindrical shape and heterogeneous structure is consistent with an organising, pre-mortem deep vein thrombus, which has embolised into the oval fossa LA - left atrium; RA - right atrium; LHIAS - Lipomatous Hypertrophy of the Inter-Atrial Septum; Thr – Thrombus; OF - Oval Fossa; * - atrial myocardium

Likewise, HiP-CT was of value in demonstrating high-contrast structures such as the 3D distribution of calcification within aortic valve, the mitral valve annulus, and coronary arteries, as well as the 3D structure of exogenous material (coronary stents) in the diseased heart (Fig. 6).

### Hierarchical anatomy from 20um to 2.2um voxel size

#### a) Atrial anatomy exemplar: Morphology of the atrial wall and of the left atrial appendage

Full-field images of both hearts at 20 µm/voxel (Fig. 3) allowed distinct anatomic characteristics of the atrial chambers to be observed, such as pectinate muscle morphology and their extent within each atrium (Fig 3a), as well as deep structure of the atrial wall in terms of endocardium, myocardium, epicardium, including fat. In both normal and abnormal heart, the endocardial layer of the left atrium was significantly thicker than the right, whereas the epicardial layer of the right atrium was significantly thicker than the left (p<0.05 for both, Fig. 3). At 6.5-2um/voxel, the histologic anatomy of the atrial walls and specifically the left atrial appendage were well-seen including lobar structure, pectinate muscle morphology, myocyte orientation, smaller atrial arterial branches as well as the clinically relevant relationship of its neck to the course of the circumflex coronary artery (Fig. 3).

**Figure 3:**
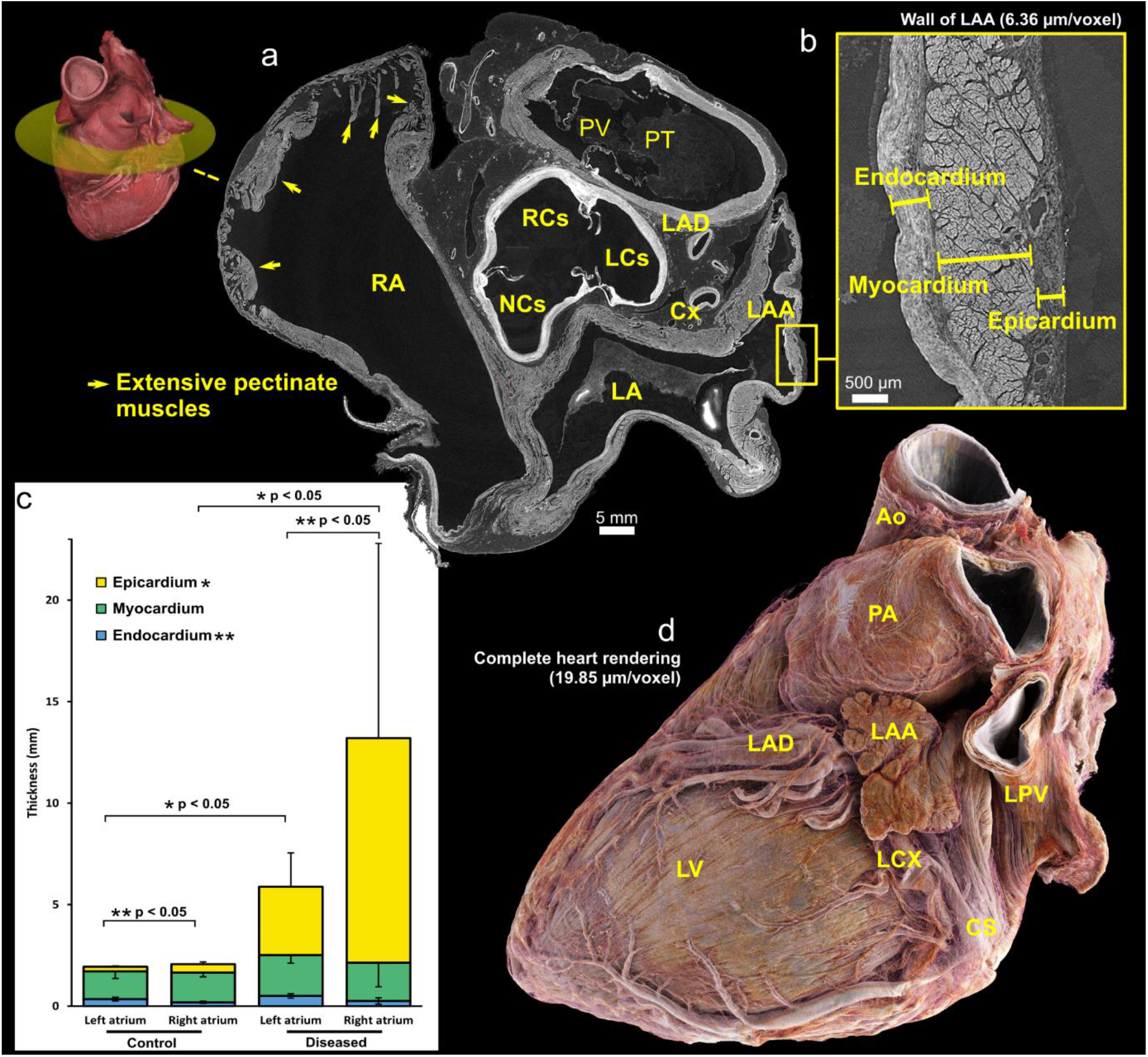
**(a)**, Short axis views of base of control heart with a 3D view indicating the location of the cross-section. **(b)** Cross-sections of high-resolution local tomography scans of the left atrial appendage wall of the control heart with a voxel size of 2.2 µm/voxel showing endocardial, myocardial and epicardial layers. **(c)**, Graph showing 2D measurements of the atrial wall layers in left and right atria for both control and diseased hearts. The thickness measurements were obtained manually at ten distinct locations along the atrial wall. Statistical analysis revealed significant differences (p < 0.05) in the thickness of the left and right atrial endocardium for both control and diseased hearts. Similarly, significant differences (p < 0.05) in the epicardial thickness between control and diseased hearts were observed for both the left and right atria which may be due primarily to extent of epicardial fat. **(d)**, 3D rendering of the control heart, with epicardial fat ‘removed’ digitally by thresholding images. The neck of the appendage can be seen running over the circumflex coronary artery – a key landmark for interventional closure of the left atrial appendage with devices used in the setting of atrial fibrillation, refractory to medical therapy for rate control, and with contra-indication to oral anticoagulants. CS – Coronary sinus; LA – Left atrium; LAA – Left atrial appendage; LAD – Left anterior descending (superior interventricular) coronary artery; LCS – Left coronary sinus; LCX – Left circumflex coronary artery; LV = Left ventricle; NCS – Non-coronary sinus; PA – Pulmonary artery; PT – Pulmonary trunk; PV – Pulmonary valve; RA – Right atrium; RCS – Right coronary sinus

#### b) Ventricular anatomy exemplars: The cardiac valves and myocardium

HiP-CT enabled visualisation of the typical components of both the atrioventricular and arterial valves (Fig. 4b, 5b), including annulus/line of attachment (respectively), valvar leaflets, tendinous cords, and papillary muscle structure. The histologic layers of the valvar leaflets (fibrosa, spongiosa, and ventricularis), could also be distinguished even in the full-field images (Fig. 5a). In the diseased heart, calcification of the aortic valve was evident predominantly within the non-coronary and the left coronary leaflets (Fig 5b). It was also possible to follow calcification within regions of the mural mitral valve annulus and mural leaflet (Fig 4b right). Mitral annular disjunction affecting the mural leaflet of the mitral valve was evident to varying degrees in both control and pathological hearts. 3D quantification demonstrated a greater severity of disjunction in the diseased heart (Supplementary Fig. 3.)

**Figure 4:**
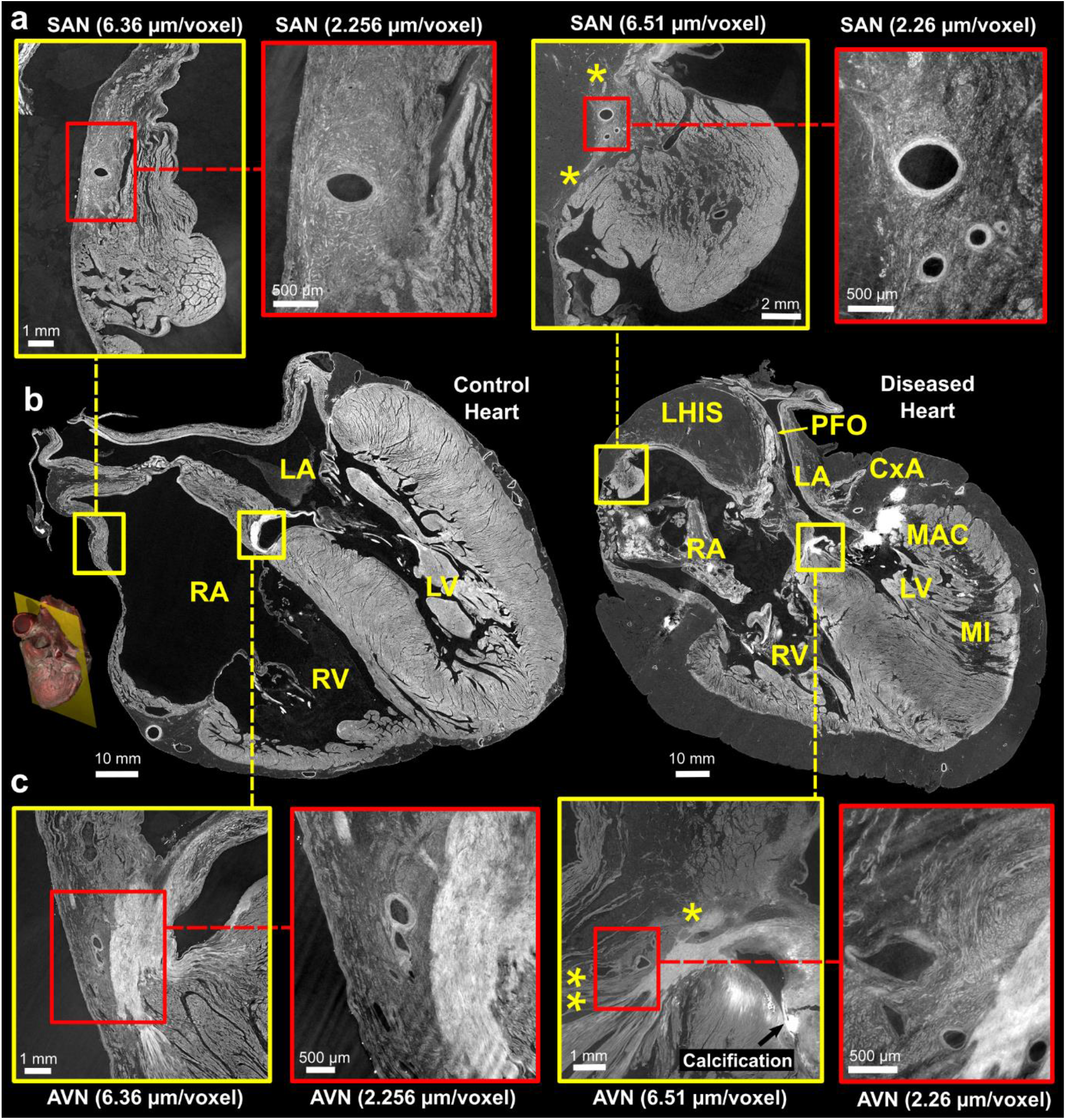
Comparison of control and pathological hearts - **(b)** Four-chamber slices of at an imaging resolution of 20 µm/voxel, with locations of the sinus node (SN) and atrioventricular nodes (AVN) highlighted. A 3D rendering provides a visualization of the imaging plane in which these views were created. Zoom images of the SAN and AVN, taken from adjacent slices, are shown in **(a)** and **(c)**, respectively. The yellow squares indicate images captured at a resolution of 6.5 µm/voxel, while the red squares indicate images captured at a higher resolution of 2.2 µm/voxel. The yellow stars in the diseased heart SAN indicate attenuated connections with working atrial myocardium running from the side of the SAN through epicardial fat and in AVN indicate connections of AVN with RA vestibule (**potential slow pathway) and attenuated connection with atrial septum (* potential fast pathway). LA - Left Atrium, RA - Right Atrium, LV - Left Ventricle, RV - Right Ventricle, MI - Myocardial Infarction, MAC - Mitral Annular Calcification, LHIS - Lipomatous Hypertrophy of the Inter-Atrial Septum, PFO - Patent Foramen Ovale. Note: The left atrium appears collapsed in both hearts.

**Figure 5:**
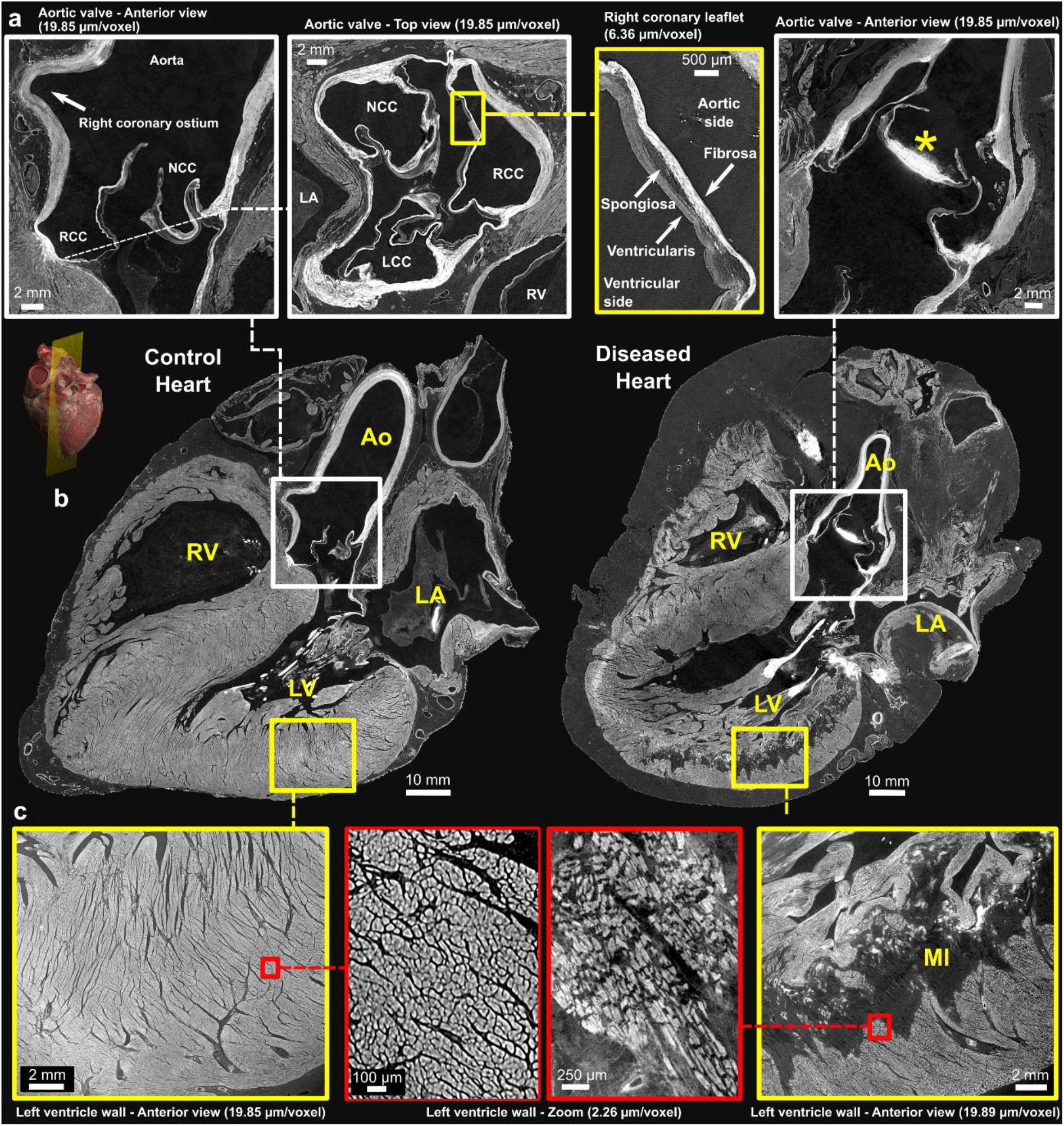
Comparison of control and pathological hearts – **(b)** Long-axis view of control and diseased hearts at imaging resolution of 20 µm/voxel. A 3D rendering provides a visualization of the imaging plane in which these views were created which is equivalent to a parasternal-long axis section. Zoomed-in images of the aortic valve and left ventricular wall are shown in **(a)** and **(c)**, respectively. The yellow squares indicate images captured at a resolution of 6.5 µm/voxel, while the red squares indicate images captured at a higher resolution of 2.2 µm/voxel. The different layers of the aortic valve, the fibrosa, spongiosa, and ventricularis, can be clearly delimited along with calcification of the non-coronary leaflet in the diseased heart (*). The arrangement of the cardiomyocytes is also clearly seen in **(c)** with altered structure and darker areas consistent with replacement fibrosis within an extensive myocardial infarction. Ao - Aorta, LA - left Atrium, RA - Right Atrium, LV - left Ventricle, RV - Right Ventricle; MI - Myocardial Infarction, NCC - Non-Coronary Cusp, RCC - Right Coronary Cusp, LCC - Left Coronary Cusp. Note: The left atrium appears collapsed in both hearts.

HiP-CT allowed for visualisation of mural architecture. In the control heart, overall myocardial architecture and orientation of myocytic aggregates, within both atrial and ventricular myocardium, is readily visible in full-field images of the whole heart, allowing for structure tensor analysis (Supplementary Fig. 4). Detail of myocytic aggregates were seen at 6.5um and individual cardiomyocytes at 2.2um (Fig 5c) but not cardiomyocyte nuclei nor intercalated discs. In the diseased heart, the posterior and inferior left ventricular free wall demonstrated a large, half thickness region of low contrast consistent with replacement fibrosis, consistent with the known myocardial infarction and coronary artery disease of the donor. At 2.2um, individual, hypertrophic cardiomyocytes can be observed surrounded by less dense areas of replacement fibrosis (Fig. 5c).

#### c) Vascular exemplar: The coronary arterial and coronary venous tree

The volumetric nature of HiP-CT data allowed for 3D visualisation of vascular anatomy without the need for further sample preparation or vascular casting. Full-field images of the whole heart allowed segmentation of the coronary arterial and venous systems down to pre-arteriolar and pre-venular level (Fig. 6d). Even small vessels, such as arteries supplying and branching from the sinus and atrioventricular nodes, as well as smaller arteries (0.25-0.75mm diameter) were clearly visible for segmentation in 20um/voxel datasets (Fig. 6c,f). The 2.2um/voxel local tomography scan enabled the observation of arterioles down to pre-capillary level including micro-vasculature within the SAN, AVN (Fig 4), and penetrating bundle of His (data not shown). The structural layers of vasculature walls in terms of tunica intima, media, and adventitia were also visible at 6.5um and below allowing differentiation between arteries and veins (Fig. 6a,b). In the diseased heart, the coronary arteries, when observed in 3D, are seen to have a distinct spiralling tortuosity, being separated from the epicardium by fatty hypertrophy and yet still anchored to the myocardium by penetrating arteries (Figs. 1, 6g).

**Figure 6:**
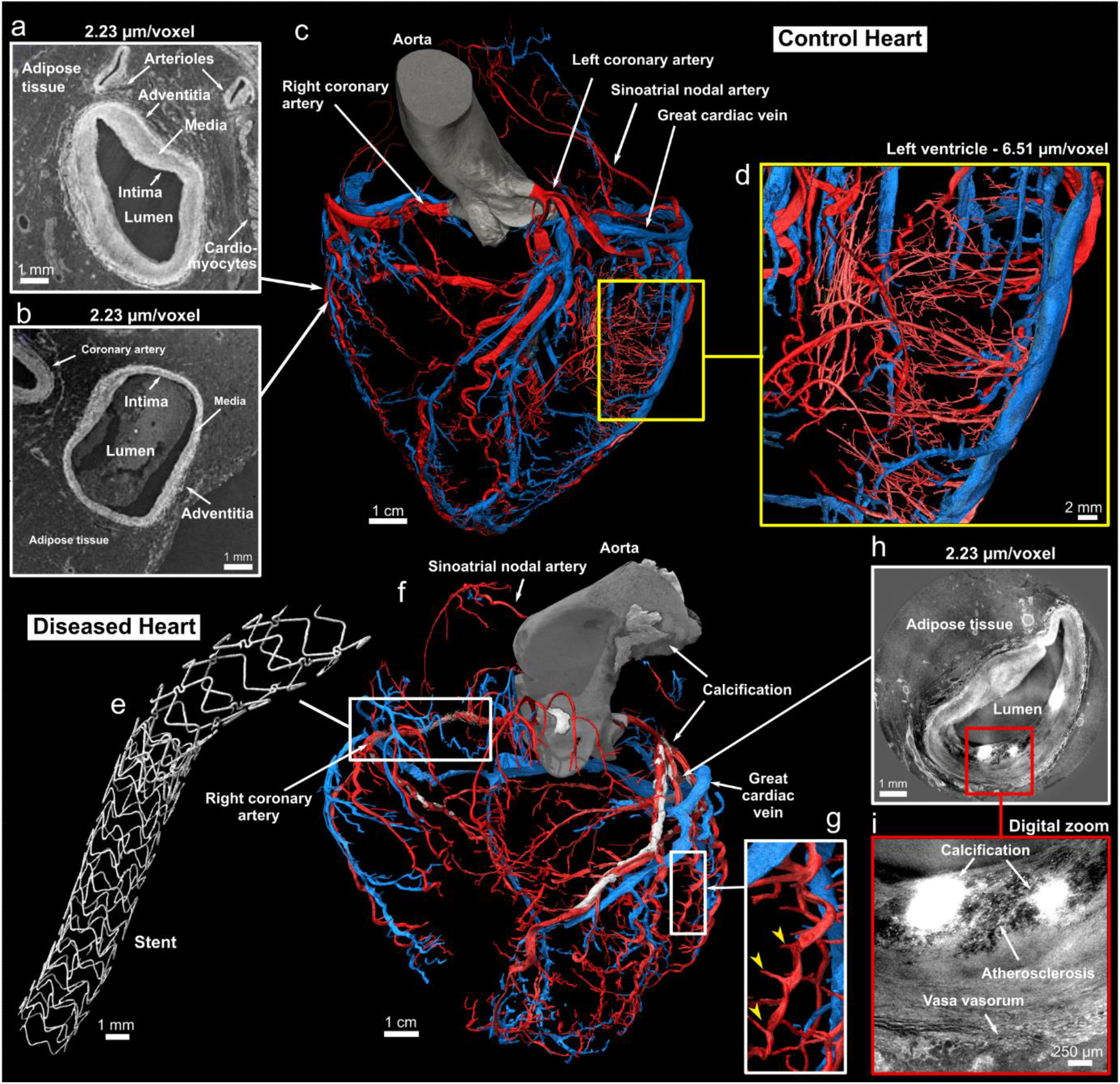
Coronary arterial and venous system of the control and diseased hearts. **(a)** and **(b)**, Cross-sections of high-resolution local tomography scans of the acute marginal artery in **(a)** and an acute marginal vein in **(b)** with a voxel size of 2.2 µm/voxel. The location of both cross-sections related to the arterial and venous coronary trees is indicated with the white arrows. Arrows indicate the layers composing the arterial and venous wall: the tunica intima, media, and adventitia. The darker tissue around the coronaries is adipose tissue, while the brighter tissue is myocytes. Segmentation of the control **(c)** and diseased **(f)** arterial (rendered in red) and venous (rendered in blue) coronary trees made from 20 µm/voxel HiP-CT images (anterior view). In **(c)**, a yellow rectangle shows a digital zoom on the left ventricular wall with segmentation of the pre-capillary arterioles in the left ventricular free wall made from a local tomography scan with 6.5 µm/voxel. Arrows indicate main coronary arteries and veins. **(d)**, Magnified view of the arteriolar segmentation in the left ventricular free wall. **(e)**, Digital zoom on one of the two stents present in **(f). (g)**, Digital zoom in **(f)** showing the marked spiral tortuosity of epicardial coronary arteries, the yellow arrows indicating how the penetrating coronary arteries act as anchor points. **(h)**, Cross-sections of a high-resolution local tomography scan of the left descending coronary artery showing atherosclerosis with calcification. **(i)**, Digital zoom in **(h)** to better show the atherosclerosis and calcification present in the arterial wall.

#### d) Exemplar of interconnectivity within the heart: The cardiac conduction system

HiP-CT was also able to demonstrate novel detail of the cardiac conduction system and its interconnectivity with surrounding structures.

### SA Node

In the control heart, the SA nodal artery arose laterally from the circumflex coronary artery and could be traced running over the roof of the left atrium, to the cavo-atrial crest, then continuing over it to descend parallel to and posterior to the terminal crest. In cross-section, the SAN was recognisable as a distinct condensation of tightly packed cells surrounding and following the course of its artery, and containing small dense, fibrous elements (Fig. 4a). Multiple longer connections between the side of the SAN and atrial myocardium/paranodal area could be traced (Fig. 4a). In long axis, the SAN cells tracked the course of the SAN artery caudally for approximately 2cm and lay immediately adjacent to and interweaved with the myocardium of the right atrium and paranodal area, throughout its length (Fig. 4a LH). SAN cells also surrounded smaller branches of the SAN artery giving the appearance of a larger volume of the SAN in these areas.

In the diseased heart, the SA nodal artery arose laterally from the stented region of the right coronary artery (RCA) although its branch point was unobstructed by the struts of the stent. It then ran towards the roof of right atrium, separated markedly from the right side of the cavo-atrial junction by (7-8mm) of epicardial fat, before encircling the junction postero-laterally, to join the SAN via a small (1mm) tortuous artery. In short axis the SAN head was small and stellate with marked infiltration of epicardial fat, which also separated the SAN along its length from the epicardium, the working myocardium, and the paranodal area within the terminal crest (Fig. 4a right). The SAN cells tracked the course of the SA nodal artery for approx 1.5cm. Connections with the adjacent atrial myocardium were still present and multiple but attenuated and stretched (Fig. 4a).

### AV Node and ventricular conduction system

The location of the compact atrioventricular node (AVN) was determined, in each heart, by following the course of the AV nodal artery (originating from the RCA), to its first branch-point (Fig. 4c). The AVN was tightly adjacent to the vestibule of the right atrium in the control heart but again separated from and infiltrated by fatty tissue on the right side in the diseased heart (Fig. 4c). Connections between the AV node and the atrial septum (potential fast pathway) and to the right and left side of the inferior pyramidal space could be traced but again were attenuated in the abnormal heart by marked fatty infiltration (Fig. 4c right).

Multiple connections between the AV node and the underlying crest of the ventricular septum could be traced in both hearts (potential nodo-ventricular pathways). Of interest, each pathway could be seen following the course of coronary arterial or venous microvasculature which carried with it varying amounts of tissue with an AV nodal phenotype, as it crossed the fibrous atrioventricular insulating tissue. In the control heart, a small potential fasiculo-ventricular loop was evident, continuous with the proximal RBB (data not shown).

The course of the penetrating bundle, (of His), and proximal left and right bundle branches (RBB) were also well-seen in both hearts (data not shown). In brief, the penetrating bundle (of His) was short with early origin of the left bundle branch in each donor. The RBB was seen as a cord running adjacent to the RV endocardium proximally towards the medial papillary muscle/cords.

## Discussion

A comprehensive understanding of the structurally normal and diseased human heart is crucial to improve understanding of pathophysiology and therefore diagnosis and treatment of CVDs. Current clinical imaging techniques, while invaluable as the mainstay for diagnosis, do not have the capacity to allow interpretation of disease across scales to cellular level. Likewise, anatomical examinations are typically either at cell level, via histology, or of gross anatomy at surgery or autopsy. There is therefore limited opportunity to understand cardiovascular disease hierarchically or trace the interconnectivity between cardiac components.

In this study, we have demonstrated the feasibility of HiP-CT to image a structurally normal and abnormal adult human heart, ex-vivo, from 20 to 2.2 µm (per voxel). We show proof of principle for mapping cardiac components in 3D at histologic resolution and for demonstrating interconnectivity between and within structures without the use of a destructive approach. This data could be linked to clinical data and or functional modelling as well as to advanced techniques such as targeted spatial transcriptomics(23).

### High resolution virtual autopsy

The study shows the utility of the technique to make high-resolution virtual autopsy diagnoses down to histologic level. As examples, we not only demonstrate the radiologic anatomy of lipomatous hypertrophy of the interatrial septum, but also entrapment of a thrombus in a patent foramen ovale. Both features are rarely seen at autopsy and difficult to demonstrate when present(29–32). Embolisation of a deep-vein thrombus, across a patent foramen ovale, which is the most likely scenario in our second donor, is seldom captured by clinical imaging, but is a clinically relevant and well-known cause of stroke, as evidenced by the increasing numbers of studies of device closure of patent foramen ovale (PFO) in this population(33). In addition, HiP-CT was able to illustrate that lipomatous hypertrophy of the ‘interatrial septum’ (LHIS), sometimes referred to as ‘lipoma of the left atrium’, is an abnormal accumulation/thickness of the epicardial fat normally found within the inter-atrial fold. As such it is sandwiched in between thin layers of right and left atrium myocardium, rather than being fatty infiltration of a ‘secondary septal’ structure.

### Potential for resolving current anatomy controversies

Multiple challenges and controversies still exist in the field of cardiac anatomy which HiP-CT has the potential to resolve. For example, comprehensive analysis of the cardiac conduction system for the interpretation of arrhythmia(34). Tracing normal and abnormal pathways or tracts between the SAN and underlying working myocardium have proven to be an arduous task and only possible by destructive serial sectioning. Our analysis shows that they have the potential for being traced in HiP-CT data, that connections surrounding the SAN are likely multiple, and are present, but attenuated, in the setting of increased adiposity. Likewise, we were able to track the course of the AVN and its vasculature supply in our datasets and show that it ran in close proximity to the overlying right vestibular myocardium and atrial septum in the control heart. This pattern, along with potential connections with the atrial septum (fast pathway), was disrupted by fatty infiltration in the diseased heart. Given the well-known association between obesity and arrhythmia(35) these findings are not only relevant to the onset of cardiac rhythm abnormalities but also to the efficacy of ablation strategies to cure them(36).

Furthermore, connections between AVN and ventricular myocardium (nodo-ventricular connections), or between the His bundle (fasciculo-ventricular) or bundle branches and the ventricular septum are notoriously challenging to document histologically, not least because sections are often lost or omitted, even with expert serial sectioning. Again, we were able to trace these connections uninterrupted in the 3D datasets. Of note, we show a novel association between such connections and the vascular supply branching from the AV nodal artery which warrants further analysis in larger numbers of hearts.

Beside arrhythmia, many anatomic questions exist which could be addressed by HiP-CT and we aimed to show potential for some of these in our study. Mapping of mural architecture within the intact ventricular mass could subsequently be correlated with advanced clinical imaging techniques, such as Diffusion Tensor Magnetic Resonance Imaging. Likewise, the 3D data allowed us to demonstrate that mitral annular disjunction was present in both control and diseased heart and show that it was more prominent in the latter, known to have mitral regurgitation. By bridging the gap between cellular and organ scales, we propose that HiP-CT, represents a step-change in radiologic techniques for resolving these and other broad-based anatomic questions relevant to the understanding and treatment of CVDs, including, but not exclusively, arrhythmia, valve, coronary and myocardial disease, as well as congenital heart disease (see Supplementary Fig. 5 for further specific examples).

## Limitations

While this study provides valuable insights, several limitations should be noted. Firstly, the imaging technique used in this study requires specialised equipment and expertise, and access to synchrotron radiation facilities is limited, which may restrict the number of specimens studied and hinder ambitious pre-clinical studies. Additionally, imaging was performed ex-vivo, which may not fully reflect the organ’s dynamic and muscular behaviour in-vivo. The major limitation preventing in-vivo scanning with HiP-CT approach is the high radiation dose associated with synchrotron imaging. Another challenge, encountered in this proof of principle study, was maintaining patency of vessels and chambers during standard whole cadaveric embalming procedures making segmentation and 3D reconstruction more difficult. To overcome this limitation, we aim to further improve sample preparation of isolated hearts in the future.

## Conclusions

This study demonstrates the remarkable potential of HiP-CT to provide high-resolution 3D images of both control and diseased human adult hearts, non-destructively and without use of contrast such as iodination. The multiscale approach enables unprecedented insights into the complex cardiac macro- and micro-structures, as well as their intricate interplay. By providing a comprehensive view of the cardiac anatomy and its alterations in disease, HiP-CT represents a promising tool for developing innovative diagnostic and treatment strategies for cardiovascular disease, ultimately leading to improved patient outcomes.

## Supporting information

Video of diseased heart - Rendered with Siemens Healthineers Cinematic Rendering

Video of control heart - Rendered with Siemens Healthineers Cinematic Rendering

## Data Availability

Complete data presented in this paper will be shared on publication on the the ESRF data repository (https://human-organ-atlas.esrf.eu).

## Code availability

The sCT data were reconstructed using a custom code written in MATLAB 2017 available on GitHub (https://github.com/HiPCTProject/Tomo_Recon) and the software package PyHST2 (https://software.pan-data.eu/software/74/pyhst2).

## Supplementary information

### Supplementary Methods

**Supplementary Figure 1:** Schematic of Hierarchical Phase-Contrast Tomography (HiP-CT) setup and data processing pipeline. Image of heart from Smart Servier Medical Art (https://smart.servier.com/) licensed under a CC BY 3.0 license. Credit for the ESRF image: ESRF, the European Synchrotron / Pixel Project.

**Supplementary Figure 2:** Visualization of the diseased heart using Neuroglancer web interface (https://github.com/google/neuroglancer). The four-pane view includes three orthogonal cross-sectional views and an independent 3-D view of the complete heart volume and all the zoom volumes, enabling to explored online the whole dataset from anywhere with a web browser and internet connection. The zoom volumes were registered to the complete heart volume semi-automatically using a non-rigid similarity transform.

**Supplementary Figure 3:** Mitral valve disjunction (MAD). Cross-section of the control (a) and diseased (b) hearts at 20 µm/voxel demonstrating the MAD in both hearts. Quantification of the MAD for the control (c) and diseased (d) hearts. A measurement of the MAD was taken every 5 degrees. MV – Mitral valve, LV - left ventricle, LA – left atrium, EF – epicardial fat, MI – Myocardial infarction, LVW – left ventricular wall, CS – coronary sinus, Cx - circumflex artery, lipomatous hypertrophy, Ao – aorta.

**Supplementary Figure 4:** Cross-section showing myocyte aggregates orientation via helical angle (HA) in the control heart at mid-ventricular level. The image is colour coded by HA values. The orientation was calculated by means of structure tensor analysis with an in-house code based on the python library structure-tensor.

**Supplementary Figure 5:** Targets areas for HiP-CT in heart. TAVI - Transcatheter Aortic Valve Implantation, TPVI - Transcatheter Pulmonary Valve Implantation, SA node - Sinoatrial Node, AV node-Atrioventricular Node, TV - Tricuspid Valve, MV - Mitral Valve, AF - Atrial Fibrillation, LA - Left Atrium, LAA - Left Atrial Appendage. Heart sketch image modified from BioRender.com.

**Supplementary Table 1:** Scan parameters for each Hierarchical Phase-Contrast Tomography scan done on both hearts. Mo = molybdenum

**Video of control heart - Rendered with Siemens Healthineers Cinematic Rendering:** movie_control-heart.mp4

**Video of diseased heart - Rendered with Siemens Healthineers Cinematic Rendering:** movie_diseased-heart.mp4

## Acknowledgements

We thank S. Bayat (INSERM) for help during the test phase, P. Masson (LADAF) for dissections of donors’ bodies, H. Dejea (ESRF) for help during the fiber orientation algorithm development, H. Reichert (ESRF) for general support of the project, and E. Boller, C. Muzelle, R. Homs, C. Jarnias, F. Cianciosi, P. Vieux, P. Cook, L. Capasso and A. Mirone for their help in setup developments and improvements. We also thank R. Engelhardt, A. Muller Brechlin, C. Petzold, N. Kroenke and M. Kuhel for help with histology and autopsies. This project has been made possible in part by CZI grant DAF2020-225394 and grant DOI https://doi.org/10.37921/331542rbsqvn from the Chan Zuckerberg Initiative DAF, an advised fund of Silicon Valley Community Foundation (funder DOI 10.13039/100014989), the ESRF funding proposals md1252 and md1290, the Royal Academy of Engineering (CiET1819/10). P.D.L., C.L.W. and JB gratefully acknowledge funding from the MRC (MR/R025673/1). P.D.L. *is* a CIFAR Fellow in the Multiscale Human Program. M.A. acknowledges grants from the National Institutes of Health (HL94567 and HL134229). AC’s research is enabled through the Noé Heart Centre Laboratories which are gratefully supported by the Rachel Charitable Trust via Great Ormond Street Hospital Children’s Charity (GOSH Charity). The Noé Heart Centre Laboratories are based in The Zayed Centre for Research into Rare Disease in Children, which was made possible thanks to Her Highness Sheikha Fatima bint Mubarak, wife of the late Sheikh Zayed bin Sultan Al Nahyan, founding father of the United Arab Emirates, as well as other generous funders.

