## Supplementary for "Multidimensional Analysis of the Adult Human Heart in Health and Disease using Hierarchical Phase-Contrast Tomography (HiP-CT)"

### Supplementary Material

#### Supplementary Methods

##### Propagation-based Synchrotron X-ray Tomography

BM18 beamline is designed to optimise propagation-based phase-contrast. The distance between the X-ray source and the sample was 150 m. We used a propagation distance of ~30 m for the full-field scan of the whole heart and between 2.5 m and 10 m for the local tomography zooms. No contrast agent was required.

The detection of the X-ray beam was performed using a LuAG:Ce 2000 µm scintillator (custom-made by Crytur, Czechia) combined with a PCO edge 4.2 CLHS (PCO Imaging, Germany) camera (2,048 × 2,048 pixels).

Two acquisition modes were used: (i) quarter acquisition[1] was used to increase the field of view to ~151 mm laterally in order to scan the whole heart, (ii) half acquisition[2] was used to performed the zooms in the heart at higher resolution. To image the whole heart or to scan large columns zooms, an automated z-series was used. Typically, for the full-field scan of the complete heart at 20 µm, the vertical field of view was ~10 mm per scan with a vertical step between scans of 7 mm, corresponding to an overlap of 30%.

For each scan of the heart in its sealed container, a scan of an equivalent container filled with ethanol 70% and agar was performed, as presented in Supplementary Fig. 1. The scan of the second container was used as beam reference for flat-field correction every successive 100 projections in order to correct most of the low-frequency effects due to local tomography.

##### Data visualization and access

Datasets were converted to N5, a format that incorporates a hierarchical structure to organize volumetric data into chunks, allowing for efficient storage and retrieval of data at different resolutions. Subsequently, the transformed datasets were transferred to the cloud, where they were made accessible through Neuroglancer[3,4], an advanced open-source viewer based on WebGL technology. Neuroglancer acts as the interface through which users can explore and interact with the volumetric data without the need for local storage or specialized hardware. In both control and diseased hearts, the 6.5 and 2.5 µm/voxel zoom volumes were registered to the 20 µm/voxel volume of the whole organ, allowing the user to visualize the different resolution at the same time (Supplementary Figure 2).

#### Analysis

##### Myocytes orientation

The local orientation of myocyte aggregates was determined using the structure tensor (ST) method. Briefly, the intensity gradients were calculated for each voxel, using the gaussian derivative method, along the three spatial orthogonal directions. Subsequently, the ST undergoes eigen-decomposition, resulting in three eigenvalues and their corresponding eigenvectors for each voxel. To identify the primary direction of the myocyte aggregates, the eigenvector associated with the smallest eigenvalue is selected. This choice is made because changes in image intensity are not expected to occur along the longitudinal axis of the cardiomyocytes. Finally, the helical angle was computed for each voxel. This angle represents the deviation between the transverse plane and the vector's projection onto the local tangential plane within the cylindrical coordinate system of the heart. The center of cylindrical system was chosen as the center of the left ventricle.

##### Mitral annular disjunction quantification

After finding the center of the left ventricle, the mitral annular disjunction was measured every 5 degrees along the mural leaflet of the mitral valve for both the control and diseased hearts.

*Supplementary Table 1: Scan parameters for each Hierarchical Phase-Contrast Tomography scan done on both hearts. Mo = molybdenum*

| Organ | Voxel size (µm) | Data label | Acquisition mode | Projection number | Propagation distance (m) | Attenuators | Average energy (keV) | Lateral field of view (mm) | Scan time (min) |
|---|---|---|---|---|---|---|---|---|---|
| Donor 1, control heart | 19.85 | Complete organ | Quarter acquisition | 9900 | 31.6 | 0.65 Mo | ~ 126 | 151.1 | 8.5 |
| | 6.36 | SAN | Half-acquisition | 6000 | 10 | 0.21 Mo | ~ 103 | 24.30 | 13 |
| | 6.36 | LAA anterior, LCA | Half-acquisition | 6000 | 10 | 0.21 Mo | ~ 103 | 24.30 | 13 |
| | 6.36 | LAA posterior, | Half-acquisition | 6000 | 10 | 0.21 Mo | ~ 103 | 24.30 | 13 |
| | 6.36 | AVN | Half-acquisition | 6000 | 10 | 0.21 Mo | ~ 103 | 24.30 | 13 |
| | 6.36 | Right ventricular myocardium wall anterior | Half-acquisition | 6000 | 10 | 0.21 Mo | ~ 103 | 24.30 | 13 |
| | 6.36 | Left ventricular myocardium free wall | Half-acquisition | 6000 | 10 | 0.21 Mo | ~ 103 | 24.30 | 13 |
| | 6.36 | Interventricular septum | Half-acquisition | 6000 | 10 | 0.21 Mo | ~ 103 | 24.30 | 13 |
| | 2.256 | SAN anterior | Half-acquisition | 6000 | 2.5 | 0.23 Mo | ~ 104 | 8.67 | 15.2 |
| | 2.256 | SAN posterior | Half-acquisition | 6000 | 2.5 | 0.23 Mo | ~ 104 | 8.67 | 15.2 |
| | 2.256 | LAD | Half-acquisition | 6000 | 2.5 | 0.23 Mo | ~ 104 | 8.67 | 15.2 |
| | 2.256 | AVN anterior | Half-acquisition | 6000 | 2.5 | 0.23 Mo | ~ 104 | 8.67 | 15.2 |
| | 2.256 | AVN middle | Half-acquisition | 6000 | 2.5 | 0.23 Mo | ~ 104 | 8.67 | 15.2 |
| | 2.256 | AVN posterior | Half-acquisition | 6000 | 2.5 | 0.23 Mo | ~ 104 | 8.67 | 15.2 |
| | 2.256 | Right ventricular myocardium wall anterior | Half-acquisition | 6000 | 2.5 | 0.23 Mo | ~ 104 | 8.67 | 15.2 |

| | | | | | | | | | |
|---|---|---|---|---|---|---|---|---|---|
| | 2.256 | Interventricular septum | Half-acquisition | 6000 | 2.5 | 0.23 Mo | ~ 104 | 8.67 | 15.2 |
| Donor 2, diseased heart | 19.89 | Complete organ | Quarter acquisition | 9900 | 31.6 | 0.65 Mo | ~ 125 | 152.08 | 8.5 |
| | 6.51 | SAN | Half-acquisition | 6000 | 10 | 0.65 Mo | ~ 113 | 25.00 | 7.1 |
| | 6.51 | RCA | Half-acquisition | 6000 | 10 | 0.65 Mo | ~ 113 | 25.00 | 7.1 |
| | 6.51 | AVN | Half-acquisition | 6000 | 10 | 0.65 Mo | ~ 113 | 25.00 | 7.1 |
| | 6.51 | Right ventricular myocardium wall anterior | Half-acquisition | 6000 | 10 | 0.65 Mo | ~ 113 | 25.00 | 7.1 |
| | 6.51 | Left ventricular myocardium free wall | Half-acquisition | 6000 | 10 | 0.65 Mo | ~ 113 | 25.00 | 7.1 |
| | 6.51 | Interventricular septum / apex | Half-acquisition | 6000 | 10 | 0.65 Mo | ~ 113 | 25.00 | 7.1 |
| | 2.26 | SAN 1 | Half-acquisition | 9900 | 2.5 | 0.44 Mo | ~ 94 | 8.66 | 11.5 |
| | 2.26 | SAN 2 | Half-acquisition | 9900 | 2.5 | 0.44 Mo | ~ 94 | 8.66 | 11.5 |
| | 2.26 | LCA branching | Half-acquisition | 9900 | 2.5 | 0.44 Mo | ~ 94 | 8.66 | 11.5 |
| | 2.26 | AVN 1 | Half-acquisition | 9900 | 2.5 | 0.44 Mo | ~ 94 | 8.66 | 11.5 |
| | 2.26 | AVN 2 | Half-acquisition | 9900 | 2.5 | 0.44 Mo | ~ 94 | 8.66 | 11.5 |
| | 2.26 | Right ventricular myocardium wall anterior | Half-acquisition | 9900 | 2.5 | 0.44 Mo | ~ 94 | 8.66 | 11.5 |
| | 2.26 | Interventricular septum | Half-acquisition | 9900 | 2.5 | 0.44 Mo | ~ 94 | 8.66 | 11.5 |
| | 2.26 | Left ventricular myocardium free wall | Half-acquisition | 9900 | 2.5 | 0.44 Mo | ~ 94 | 8.66 | 11.5 |

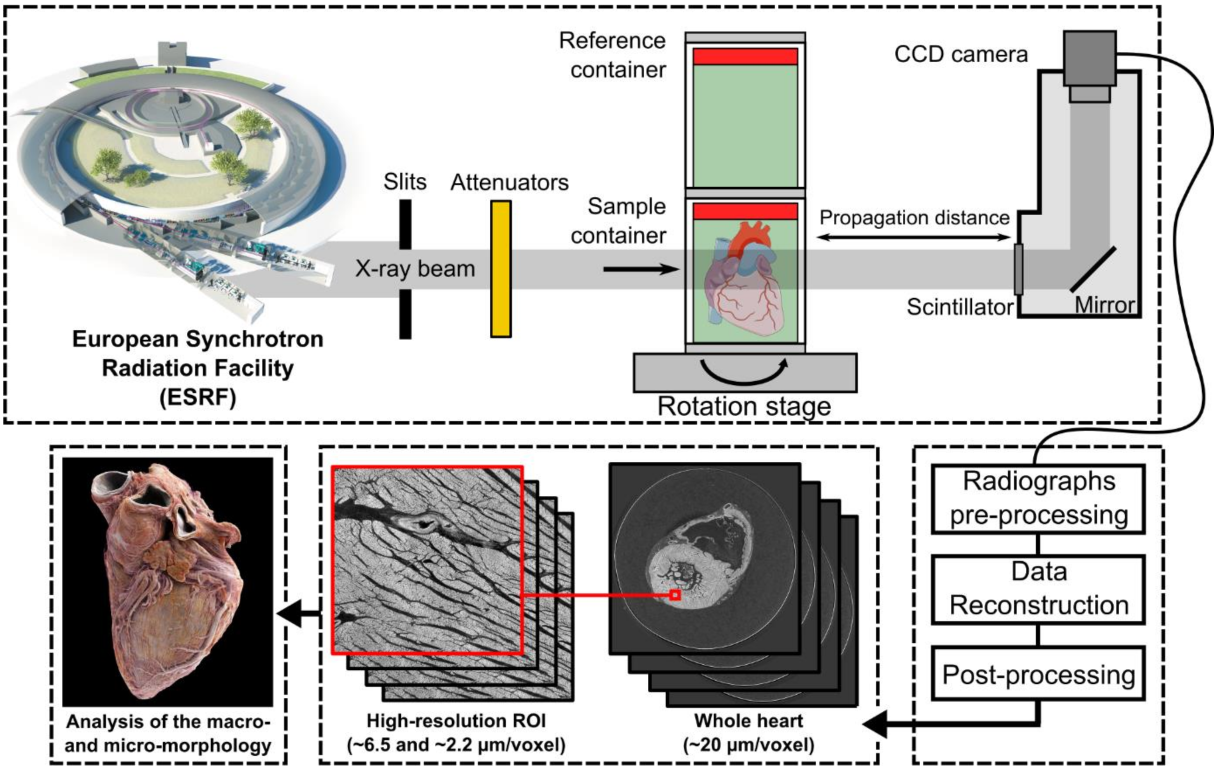


*Supplementary Figure 1: Schematic of Hierarchical Phase-Contrast Tomography (HiP-CT) setup and data processing pipeline. Image of heart from Smart Servier Medical Art (https://smart.servier.com/) licensed under a CC BY 3.0 license. Credit for the ESRF image: ESRF, the European Synchrotron / Pixel Project.*

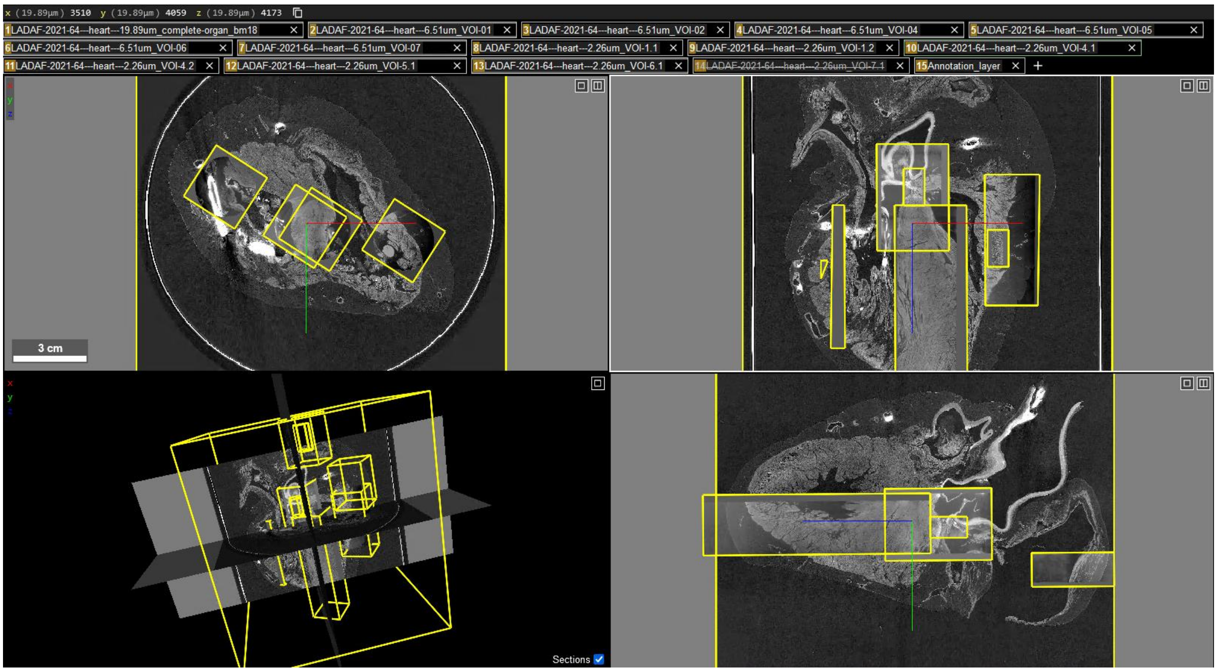


*Supplementary Figure 2: Visualization of the diseased heart using Neuroglancer web interface (https://github.com/google/neuroglancer)[3,4]. The four-pane view includes three orthogonal cross-sectional views and an independent 3-D view of the complete heart volume and all the zoom volumes, enabling to explored online the whole dataset from anywhere with a web browser and internet connection. The zoom volumes were registered to the complete heart volume semi-automatically using a non-rigid similarity transform.*

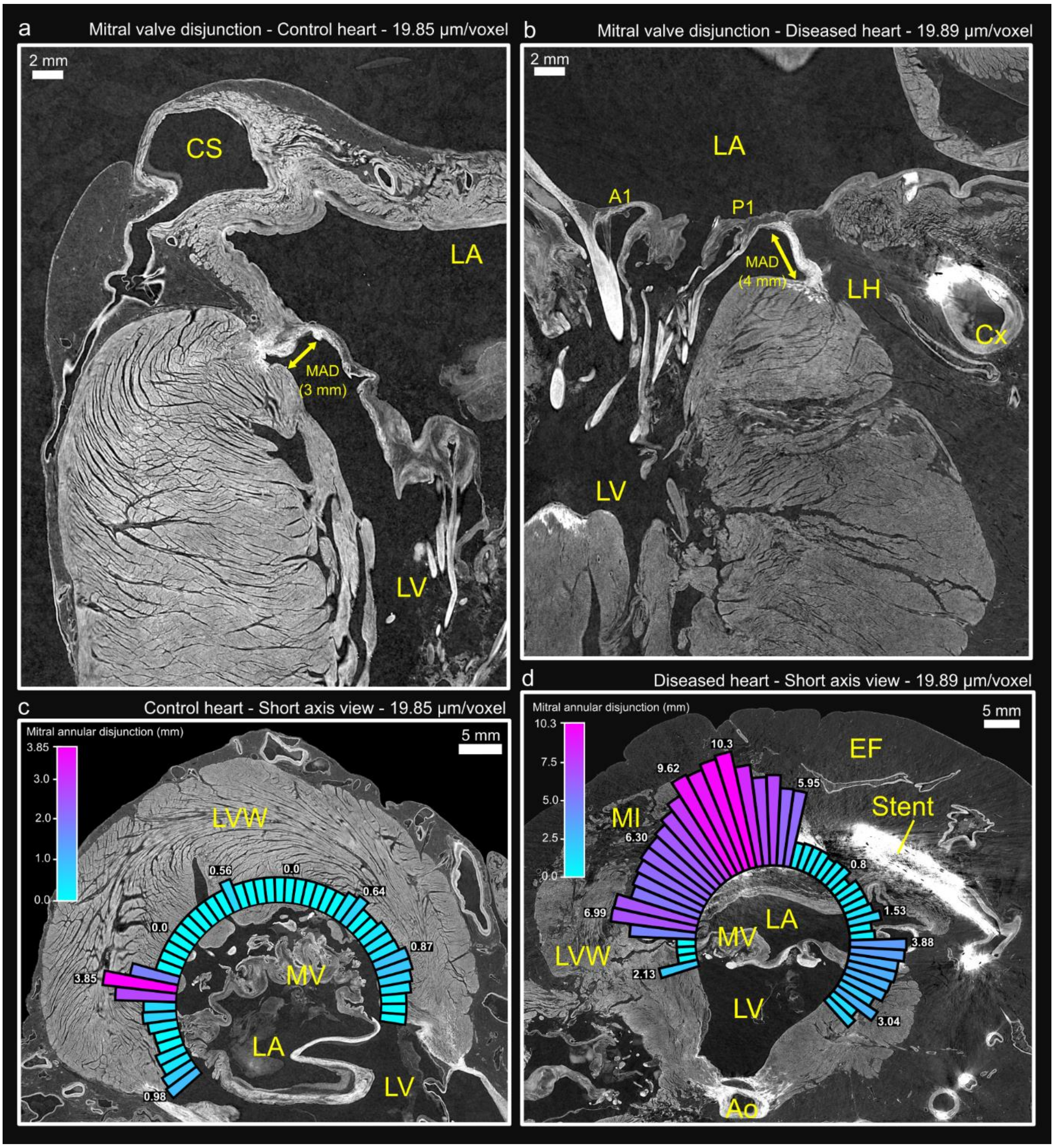


*Supplementary Figure 3: Mitral valve disjunction (MAD). Cross-section of the control (a) and diseased (b) hearts at ~20 µm/voxel demonstrating the MAD in both hearts. Quantification of the MAD for the control (c) and diseased (d) hearts. A measurement of the MAD was taken every 5 degrees around the mural leaflet. MV – Mitral valve, LV - left ventricle, LA – left atrium, EF – epicardial fat, MI – Myocardial infarction, LVW – left ventricular wall, CS – coronary sinus, Cx - circumflex artery, lipomatous hypertrophy, Ao – aorta.*

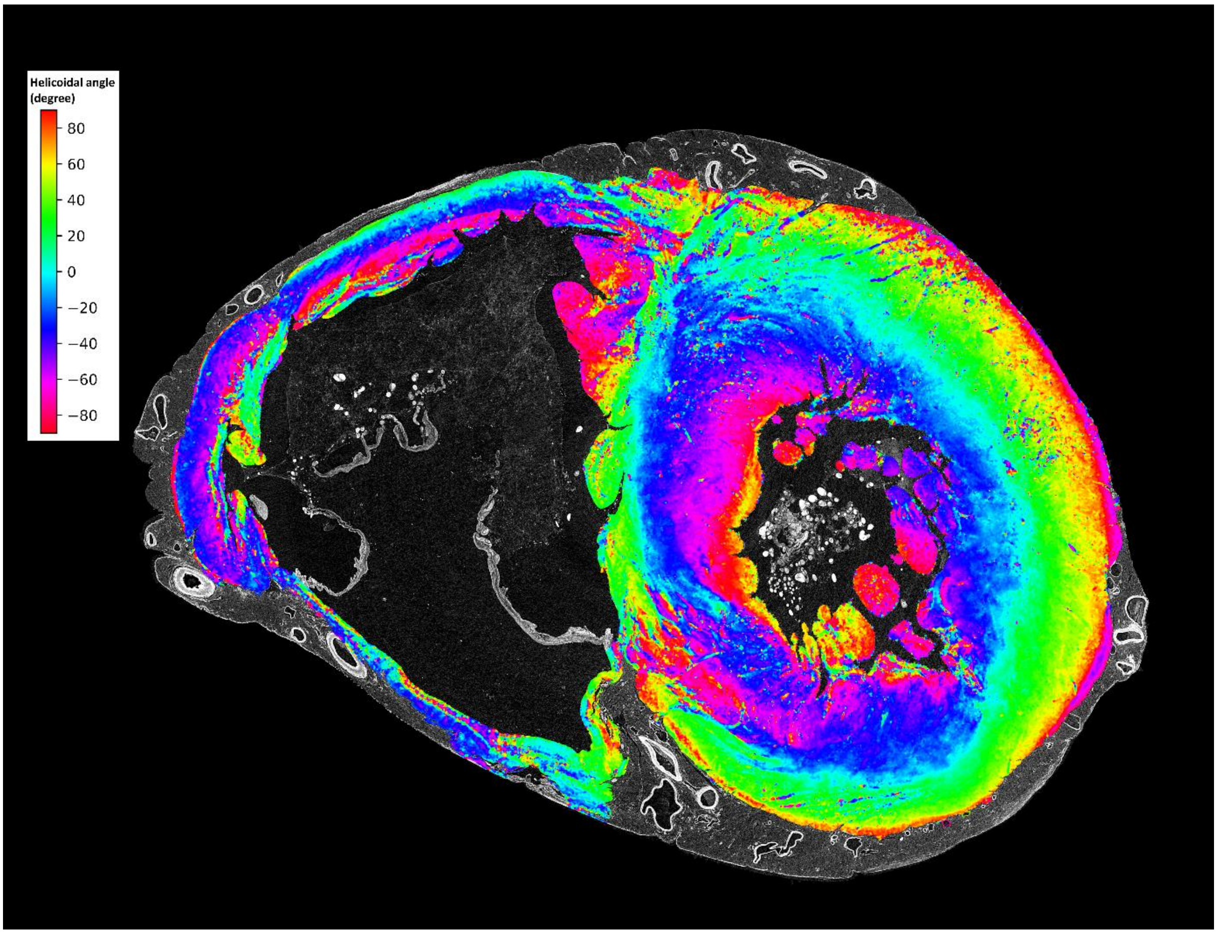


*Supplementary Figure 4: Cross-section showing myocyte aggregates orientation via helical angle (HA) in the control heart at mid-ventricular level. The image is colour coded by HA values. The orientation was calculated by means of structure tensor analysis with an in-house code based on the python library structure-tensor[5,6].*

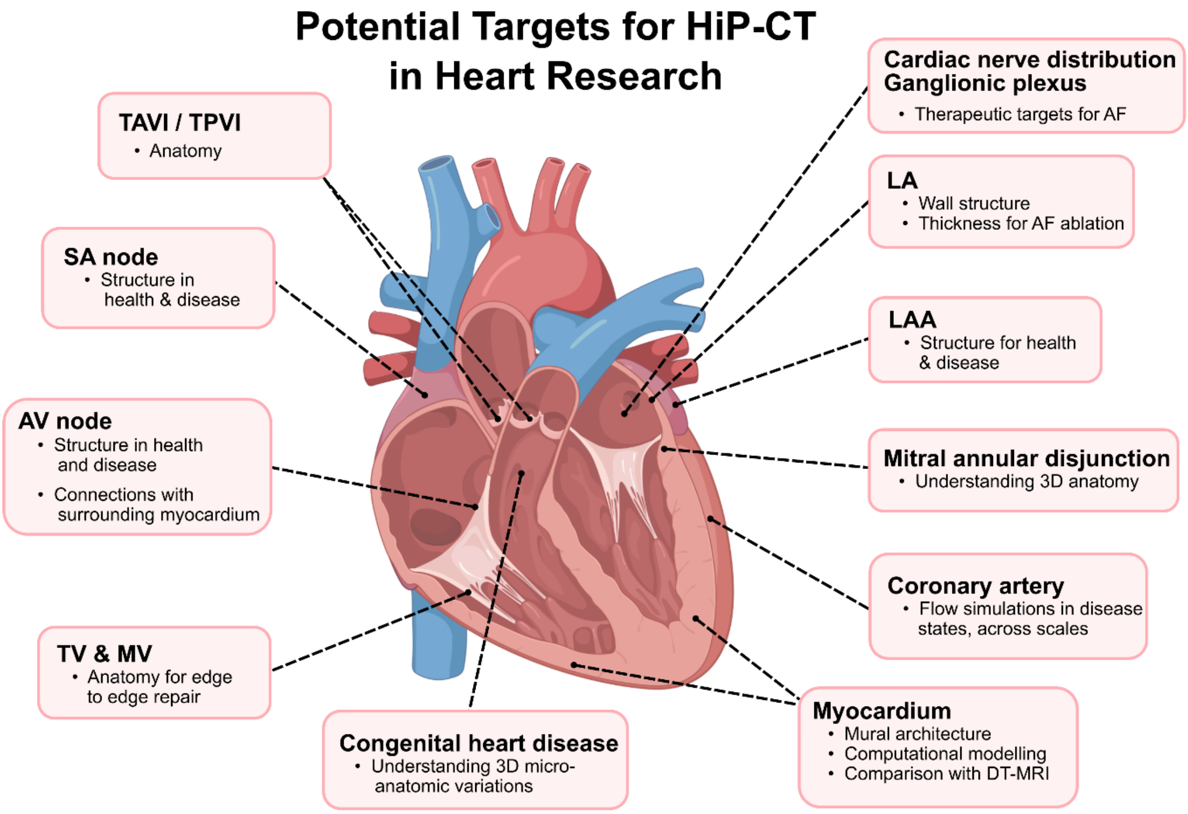


*Supplementary Figure 5: Targets areas for HiP-CT in heart. TAVI - Transcatheter Aortic Valve Implantation, TPVI - Transcatheter Pulmonary Valve Implantation, SA node - Sinoatrial Node, AV node - Atrioventricular Node, TV - Tricuspid Valve, MV - Mitral Valve, AF - Atrial Fibrillation, LA - Left Atrium, LAA - Left Atrial Appendage. Heart sketch image modified from BioRender.com.*
